# Impact of the pulse artifact on evoked activity in human wakefulness and sleep

**DOI:** 10.1101/2025.11.20.689416

**Authors:** Jacinthe Cataldi, Andria Pelentritou, Sophie Schwartz, Marzia De Lucia

## Abstract

The investigation of the neural evoked response to the heartbeat quantified using electroencephalography (i.e., heartbeat evoked potentials or HEPs), has gained recent attention in neuroscience, notably as a measure of interoception, the sensory system responding to internal bodily states. One main challenge in measuring HEPs is their susceptibility to cardiac artifacts contamination, including the cardiac field artifact and the pulse artifact (PA), the latter possibly caused by the mechanical pressure of pulsating vessels. Here, we aimed at assessing the impact of PAs on HEPs and auditory evoked potentials (AEPs, a proxy of the neural responses to exteroceptive sensory stimuli). To this aim, we compared two pre-processing pipelines using independent component analysis in healthy volunteers (N=30). The first, standard, pipeline excluded ocular, muscle, sweating-related activity and cardiac-related activity stemming from the cardiac field artifact. The second, pairwise phase consistency (PPC) pipeline, in addition to the removal of the aforementioned components, used the quantitative metric of PPC between independent components and the ECG to remove the cardiac-related PA. We tested how these two pre-processing approaches influenced HEPs and AEPs recorded under diverse neurophysiological conditions (wakefulness, N2, N3, and REM sleep). Comparing the HEPs from the standard and the PPC approaches (cluster-based permutation statistics, p<0.05, two-tailed) revealed a significant effect of PAs, particularly in wakefulness, followed by REM and N2 sleep, with Cohen’s d effect sizes of 1.92, 0.95 and 0.88, respectively. By contrast, PA correction had a negligible effect (p>0.05, two-tailed) on the HEPs in N3 sleep and on the AEPs in all vigilance states. Our results emphasize the need to account for the PA as a significant confounding factor when comparing HEPs across groups with varying vascular or cardiac conditions.

## 1. Introduction

Our brain continuously receives signals from the body to monitor the internal physiological state. The ability to perceive and process these internal signals is known as interoception, and it plays a fundamental role in maintaining physiological homeostasis and shaping emotion, cognition, and behavior [1–3]. Among these internal signals, the heartbeat provides a rhythmic stream of afferent inputs to the brain, making it a key channel through which interoception can be studied. Specifically, the heartbeat evoked potential (HEP), an electroencephalographic (EEG) event-related potential time-locked to the R-peak of the electrocardiogram (ECG) [4–6], is thought to reflect the neural response to visceral afferent inputs from baroreceptors and cardiac mechanoreceptors, conveyed via the vagus nerve to subcortical and cortical regions, including the insula, anterior cingulate, somatosensory, and prefrontal cortices [7–10]. Notably, the HEP can inform about interoceptive processing during explicit interoceptive tasks as well as during resting wakefulness, sleep [8, 11–13], and disorders of consciousness [14, 15]. Sleep is of particular interest as it provides a natural model of varying degrees of sensory disconnection and neural dynamics, with distinct vigilance states characterized by different background EEG activity and bodily signals’ dynamics. Indeed, some studies have documented a modulation of HEP amplitude across sleep stages [8, 13, 16]. Specifically, HEPs recorded during N2 and N3 sleep display large differences in amplitude compared to rapid eye movement (REM) and stage 1 of non-REM (N1) sleep, the latter two more closely resembling those observed during wakefulness [8, 16].

Despite increasing interest in the HEP, its reliable measurement is technically challenging due to the contamination of the EEG signal by cardiac-related artifacts. By construction, HEPs are temporally locked to the heartbeat and thus overlap with at least two known and distinct types of cardiac artifacts recorded by scalp electrodes: the electromagnetic field generated by cardiac electrical activity and the mechanical effects of cardiac-induced vascular pulsations [17–19]. The electromagnetic cardiac field artifact (CFA) has been the focus of most cardiac artifact removal approaches, and it is typically characterized by a high voltage deflection time-locked to specific phases of the cardiac cycle, primarily the QRS complex and T wave [20]. Topographically, this artifact is characterized by a left-right asymmetric voltage distribution with equipotential lines running diagonally from the left front to the right back (Figure 1B) [20].

**Figure 1.**
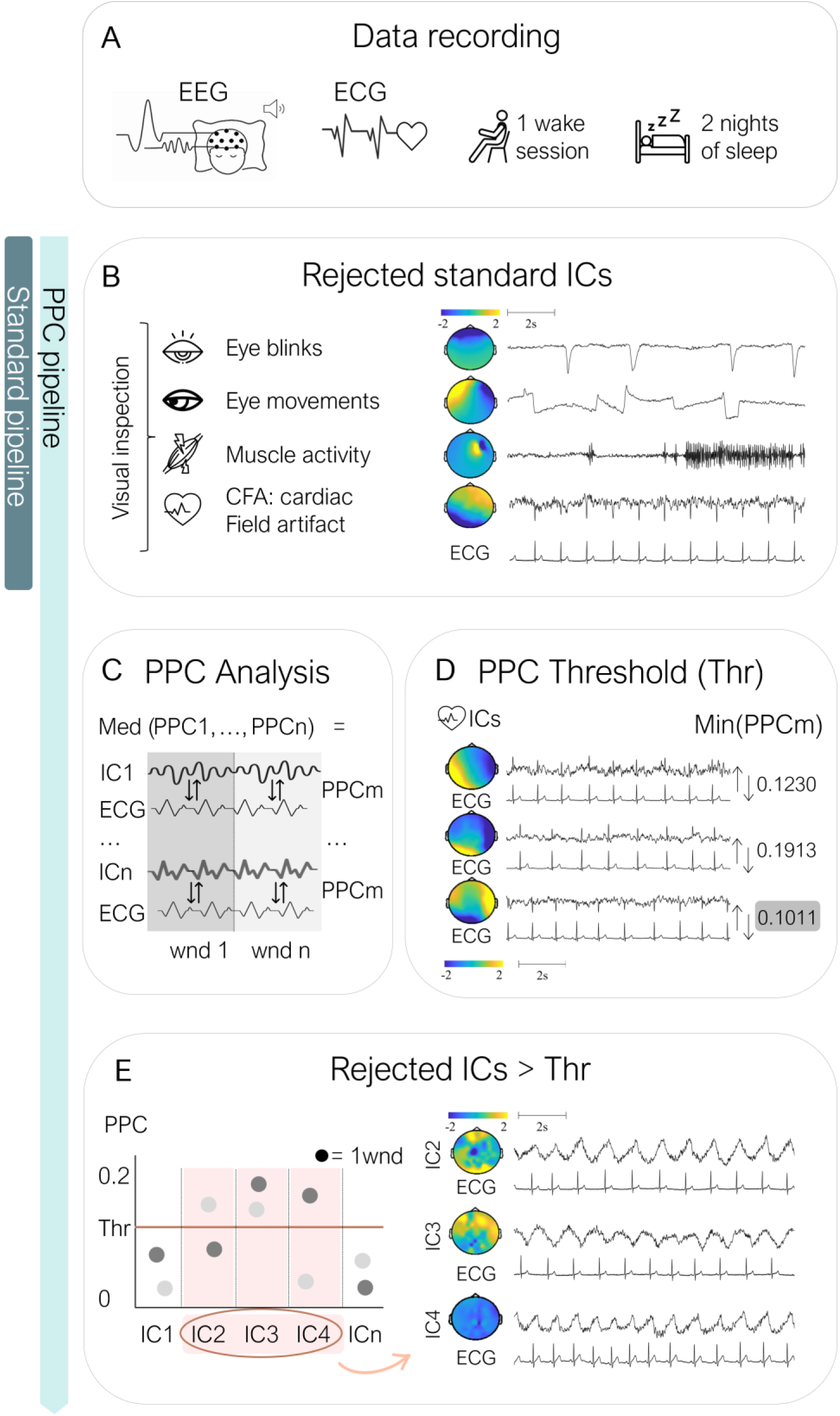
Schematic overview of the standard and cardiac pre-processing pipeline for EEG data. EEG and ECG data were recorded during one wakefulness session and two overnight sleep sessions. The standard pre-processing pipeline included the visual inspection and rejection of independent components (ICs) associated with artifacts related to eye blinks, eye movements, muscle activity and cardiac field artifacts. The pairwise phase consistency (PPC) pipeline considered all the ICs identified in the standard pipeline and additionally considered ICs related to pulse artifacts. PPC was computed between each IC and the ECG signal across time windows (wnd), and the median PPC value (PPCm) was derived for each IC across non-overlapping windows of 1000 R-peaks. A rejection threshold (Thr) was defined as the minimum PPCm across all ICs. ICs exhibiting PPC values exceeding this threshold were considered cardiac-related and removed from further analyses after visual inspection. Topographical maps and time series examples illustrate ICs retained or rejected at each step of the two pipelines.

Differently from the CFA, which reflects a fixed electromagnetic field pattern, the pulse artifact (PA) is related to the pulsatile blood flow and may exhibit considerable variability across individuals as it is inherently linked to the subject-specific vascular anatomy [21]. Furthermore, the PA strength is related to the blood supply, which may vary within subjects depending on the spatial distribution of activated neural sources [19]. A higher blood supply can lead to stronger pressure on the EEG electrodes and consequent variation in the impedance level [19]. Because PAs depend on the ‘mobility’ of single electrodes, they also vary as a function of the participant’s head shape and the EEG cap tightness. Finally, the PA also partly reflects the small head and body movements due to the heart contractions and blood ejection [22]. The electrical signature of the PA has been described as a lateralized or diffuse rhythmic slow wave following the QRS complex with amplitude modulation at approximately 200 ms after the R peak (Figure 1E) [21–26].

The PA is well recognized and extensively addressed in simultaneous EEG-fMRI recordings where such movements are of high interest since interactions between the cardiac pulse and the magnetic field can result in signal distortion [21, 27]. Similarly, in intracranial recordings [4], signals have been shown to exhibit PAs. Specifically in electrocorticography, PA amplitudes can be up to six times larger than those observed in simultaneously recorded scalp EEG, likely due to the closer proximity of electrodes to cerebrovascular and mechanical sources [28]. However, the impact of PAs is not explicitly addressed in most EEG pre-processing pipelines or conflated with the CFA [29]. The present study thus aims at assessing the potential of PAs to significantly contaminate HEPs in scalp EEG recordings.

While most previous studies on the impact of cardiac related artifacts on scalp EEG were conducted on healthy awake individuals [19, 29, 30], the extent of the EEG contamination by cardiac related artifacts across different vigilance states remains largely unexplored. In this context, it is plausible that the EEG during wakefulness and sleep will be contaminated to a variable degree due to differences in EEG background activity in amplitude and spectral composition [31, 32]. In addition, autonomic nervous system activity undergoes systematic changes along the sleep-wake cycle, with increased sympathetic activity during wakefulness and REM sleep, and predominantly parasympathetic activity during non-REM (NREM) sleep, directly affecting both cardiac activity and the HEP [33–39]. These state-dependent factors may shape the relative contribution of cardiac artifacts on EEG signals, raising the need for the evaluation of their impact across vigilance states.

Here, we investigated the impact of cardiac related artifacts, and particularly the PA, in the same healthy volunteers during wakefulness and sleep. First, we hypothesized that cardiac artifacts are more prominent when the EEG background has low amplitude, consistent with previous reports of PAs during low-amplitude states such as REM sleep [24]. By contrast, during N3 sleep, the presence of high-amplitude oscillations, such as slow waves, may partly mask or outweigh cardiac contributions [40]. Second, since the PA may primarily reflect the mechanical transmission of blood through cranial vessels, its influence should increase with higher blood pressure. Blood pressure is known to be higher in wakefulness and REM sleep and attenuated in NREM sleep [35], with an additional increase expected during wake recordings in a seated position. Based on these points, we hypothesized that both the CFA and PA would be more common in wakefulness and REM sleep compared to NREM sleep, with N3 showing the lowest level of contamination.

Understanding these differences is of relevance for the computation and correct interpretation of HEPs as well as for informing the choice of cardiac artifact correction methods. In this work, we propose a semi-automated method combining Independent Component Analysis (ICA) with pairwise phase consistency (PPC) to improve the detection of cardiac-related independent components (ICs). PPC [41] quantifies the consistency of a phase relationship between the time-course of each IC and the ECG signal, making it particularly effective at detecting both the CFA and the PA regardless of their amplitude characteristics [19]. This phase-based approach is advantageous because both types of cardiac artifacts have an inherent temporal relationship with the heartbeat, even when their amplitude correlation to the ECG may be weak. We compared this approach to a more traditional ICA-only pipeline that relies on visual inspection alone to reject ICs showing CFA. Our first aim was to assess the impact of these pre-processing strategies on EEG signals by applying each of them to the same dataset and comparing their effects on HEPs across vigilance states. We additionally analyzed auditory-evoked potentials (AEPs) in response to sounds presented without any temporal relationship to the ongoing heartbeat. Finally, we used this methodological framework to better describe the prevalence of cardiac artifacts across vigilance states, especially for the under-investigated PA. In doing so, we aimed at providing novel insights into how the PA contaminates HEPs and non-cardiac related evoked activity i.e., AEPs.

## 2. Methods

### 2.1. Ethics statement

The study (Project-ID: 2020-02373) was approved by the local ethics committee (La Commission Cantonale d’Ethique de la Recherche sur l’Etre Humain), in accordance with the Helsinki declaration.

### 2.2. Participants

Thirty-three healthy volunteers (mean±standard deviation age = 23.42±3.48 years, 16 female) with good sleep quality enrolled in the study. Eligibility was determined through a structured phone interview, which verified the absence of hearing issues or psychiatric, neurological, respiratory, and cardiovascular disorders, including sleep apnea. Additional inclusion criteria included a regular sleep–wake schedule without excessive daytime sleepiness or sleep disturbances, verified using the Pittsburgh Sleep Quality Index and the Epworth Sleepiness Scale, respectively. All participants provided written informed consent and received approximately 420 Swiss Francs as compensation for their participation.

### 2.3. Experimental Procedures

The study employed a two-way crossover design, with participants undergoing three consecutive overnight sleep sessions and a separate wakefulness session. The wakefulness session was scheduled either within one week before the first sleep session or within one week following the final sleep session; for 15 participants the wakefulness session was conducted first.

The wakefulness session was conducted in a dimly lit experimental booth isolated from sound and electrical interference. Participants fixated on a centrally located white cross displayed on a grey screen with their head position maintained using the eye-tracker chin rest, placed approximately 55 cm from the screen. The sleep sessions were conducted on consecutive nights in a bedroom with sound and electrical insulation. Participants filled out a standardized sleep and dream diary for eight days prior to the first sleep session and in the mornings following the three nights at the laboratory to monitor their sleep habits and dream content. In addition, actigraphy monitoring (Actigraph GT3X+, ActiGraph, FL, USA) for three days before the first sleep session ensured a regular sleep schedule prior to sleeping at the laboratory. The first sleep session was an adaptation night during which 6-lead EEG, horizontal and vertical electrooculography (EOG), and submental electromyography (EMG) were acquired using Ag/Au cup electrodes and the v-AMP amplifier (Brain Products GmbH, Gilching, Germany). During adaptation, a sequence of 100 stimuli (pure tones, 400 Hz) was presented binaurally using earphones every hour of sleep to ensure participants could sleep efficiently upon auditory stimulation. Sleep scoring of the adaptation data verified a high sleep efficiency, after which participants were asked to return to the laboratory for the two experimental sessions that included the auditory stimulation protocol.

### 2.4. Data acquisition

During the wakefulness session and two experimental nights, EEG signals were acquired using the 64-channel ANT Neuro EEG system (eego mylab, ANT Neuro, Hengelo, Netherlands) sampled at 1024 Hz and lab streaming layer (LSL, https://github.com/sccn/labstreaminglayer), with electrode CPz serving as the reference and AFz as the ground. Electrode impedance was set at 5 kΩ. Concomitant to the EEG, we acquired ECG signals through a two-lead setup with two electrodes placed on the chest (one on the right infraclavicular fossa, one below the heart). Vertical and horizontal EOG were acquired using electrodes placed below the left eye and next to the left and right canthi, respectively. Submental EMG and respiration using a belt placed around the diaphragm were additionally recorded. Further, during the wakefulness session alone, ocular data were acquired at 500 Hz using the SR Eyelink 1000 plus (SR Research Ltd., Mississauga, Canada). Auditory stimuli were presented binaurally through in-earphones.

### 2.5. Auditory stimulation

During wakefulness and the two experimental nights of sleep, Psychopy [42] and custom-made Python scripts were used for paradigm administration. The experimental paradigm consisted of the administration of three types of auditory sequences as extensively described elsewhere [43]. and a baseline condition without auditory stimulation. In this study, we focused only on the condition where sounds occurred at fixed time intervals, commonly utilized in auditory regularity processing studies [44]. However, all pre-processing steps outlined below were performed including data from all experimental conditions. During wakefulness and sleep, experimental conditions were presented in blocks, each comprised by the baseline lasting 4 minutes, and the auditory conditions (300 stimuli each), presented in a pseudo-random order. All conditions within a block were separated by an interval of 30 seconds and the experimental block of four conditions lasted approximately 20 minutes (depending on the individual’s heart rate). A pause for participants to rest occurred after each block of four conditions in wakefulness until a total of six blocks was administered while, in sleep, paradigm administration ensued without pauses.

Sounds were 1000 Hz sinusoidal 16-bit stereo tones of 100 ms duration and 0 μs inter-aural time difference sampled at 44.1 kHz and generated using the *sound* function in Psychopy [42]. A Tukey window was applied to each tone. Sound stimuli were presented binaurally through in-earphones with an intensity adjusted for each participant to a comfortable level, determined prior to the experimental paradigm administration. Sound intensity was different between wakefulness and the two acquisition sleep sessions to avoid perturbing sleep by high volume auditory stimuli.

### 2.6. EEG and ECG data analysis

Data were analyzed using MATLAB (R2019b, The MathWorks, Natick, MA) using the FieldTrip [45] and EEGLAB [46] toolboxes, and custom scripts.

#### 2.6.1. Pre-processing

The sleep electrophysiological data acquired over two experimental nights were scored by an expert sleep technician (SOMNOX SRL, Belgium) who was blind to the experimental manipulation in this study, following the AASM guidelines for sleep scoring [47]. Continuous EEG and ECG data from wakefulness and sleep recordings were bandpass filtered between 0.5 and 30 Hz. Sleep data was segmented into N2, N3 and REM, excluding portions of data scored as N1 sleep, ‘wake after sleep onset’ or arousals.

For each participant and session (wakefulness, each of the two experimental nights), two distinct pre-processing pipelines were applied to the EEG data, separately for each vigilance state (Wake, N2, N3, REM) resulting in two pre-processed datasets (Figure 1). R-peaks in the ECG signal were detected for all vigilance states using the Pan-Tompkins algorithm [48]. Experimental blocks with frequent flawed real-time R peak detection or faulty auditory stimulus presentation were excluded.

#### 2.6.2. Standard pre-processing

EEG data was visually inspected to identify and exclude periods with excessive noise across electrodes. Artifact electrodes were identified and excluded using a semi-automated approach implemented in FieldTrip [45], as well as the visual inspection of individual electrodes throughout the recording periods (wake: 7.4 ± 2.4, mean ± SD; N2: 13.7 ± 3.2; N3: 7.3 ± 2.1; REM: 11.5 ± 2.8). Next, ICA was performed separately for each participant during each vigilance state and night using the Infomax algorithm (runica), as implemented in EEGLAB [46], to identify and remove ocular, muscle, sweating, and cardiac related activity.

The identification and removal of cardiac related ICs exhibiting CFAs was based on the visual inspection of the ICs topographic and temporal features using the following criteria: (1) a characteristic left–right asymmetric topography, (2) the presence of a sharp deflection concomitant to the ECG R peaks throughout the recording, assessed through the visual inspection of the IC time course alongside the ECG signal (Figure 1B).

#### 2.6.3. PPC pre-processing

In addition to the pre-processing steps described above for the standard pipeline, in a separate pre-processing analysis, we aimed at selecting components related to the PA within each vigilance state. These components were identified and removed through a quantitative analysis of the phase consistency between each IC and the ECG signal based on pairwise phase consistency (PPC), as implemented in FieldTrip [19, 45]. In this study, we considered the PPC in the 0-25 Hz range, since we expected that EEG activity related to the PA would not spam frequencies above 25 Hz [19]. To allow for the accurate identification of PA-related components, EEG and ECG data were segmented into time-windows containing 1000 R peaks. This time-window length was chosen in order to match the shortest period of N3 and REM sleep in our dataset, allowing for the retainment of 95% of data in each vigilance state and participant. On average, this procedure yielded 7.5 ± 5 (mean ± standard deviation) windows during wakefulness, 11.7 ± 3.4 during N2 sleep, 3.4 ± 1.3 during N3 sleep, and 5 ± 1.8 during REM sleep. For a given time-window, we computed the PPC between each IC and the ECG signal in non-overlapping epochs extracted from −100 ms to 500 ms relative to the R peaks of the ECG, to focus on the period likely to contain cardiac-related artifacts in the EEG (differently from [19] that utilized −200 ms to 200 ms epochs). These analyses resulted in one PPC value ranging from 0 to 1 for each time-window of 1000 non-overlapping epochs (Figure 1C).

To establish a PPC threshold value for classifying ICs as cardiac related, we took as reference the PPC value obtained from the ICs visually identified as CFA in the standard pre-processing pipeline. The median PPC value was computed across time windows for each CFA component for each participant and for each vigilance state (Wake, NREM (N2 and N3 combined), REM) in which the CFA was identified as part of the standard pre-processing. Next, within each vigilance state (Wake, NREM, REM) and across the two experimental nights, the minimum of these median PPC values from CFA components across subjects and experimental night was adopted as the threshold value. This conservative approach ensured that only components with phase-locking strength at least equal to the weakest CFA would be selected for further inspection (Figure 1D). ICs exceeding this threshold in at least one time-window were visually inspected. Components were classified as contaminated by cardiac artifacts through visual inspection if they showed time-locked activity to the ECG signal with PA typical morphology, characterized by broader, slower waveforms [28]. The inspection focused specifically on time windows where the PPC exceeded the threshold, ensuring that phase-locking and morphological criteria were evaluated in parallel (Figure 1E). We also identified and removed components related to the CFA that were overlooked in the standard pre-processing pipeline. Components exhibiting cardiac-related artifacts in at least one time-window of 1000 epochs were subsequently rejected for the entire vigilance state’s recording.

The EEG signal was reconstructed following the removal of the artifact-related components separately for the standard and PPC pre-processing pipeline. Finally, we interpolated artifact electrodes using the spline method [49].

#### 2.6.4. Evoked activity

The two datasets, resulting from the two pre-processing pipelines, were segmented from –100 ms to +600 ms relative to R-peaks in the baseline to compute HEPs, and relative to auditory stimuli in the isochronous condition to compute AEPs. Trials in which the analysis window overlapped with neighboring events (e.g., due to short RR intervals <700 ms) were excluded. A semi-automated trial rejection procedure was performed separately for HEPs and AEPs using FieldTrip [45], for each participant, vigilance state and night. Trials were visually inspected and rejected when identified as outliers based on absolute amplitude, standard deviation, kurtosis, z-score, and variance. Trial rejection was performed separately for each dataset resulting from the two pre-processing pipelines and only trials retained in both datasets were further considered in subsequent analyses. Finally, EEG data were re-referenced to the common average.

For comparisons between pre-processing pipelines, HEPs and AEPs were computed for each subject and vigilance state following concatenation of the datasets from the two nights for sleep. Next, an average using an equal number of trials from each dataset was computed. Importantly, the selected trials for the standard and PPC pipeline were non-overlapping, ensuring that each pipeline was evaluated on an independent subset of the data. Subjects with fewer than 100 trials for a given vigilance state were excluded from the corresponding analysis. In addition to the evoked activity, we computed the Global Field Power (GFP) by calculating the standard deviation of the single-trial evoked activity across channels at each time point on the subject-averaged evoked response, separately for the two datasets obtained from the two pre-processing pipelines. We then averaged the resulting single-trial GFP values within each dataset across subjects. The GFP was computed to quantify the strength of the evoked activity at each time point within the –100 ms to 600 ms trial, relative to the event of interest. We expected to observe equal or higher average GFP values in the dataset processed with the standard compared to the PPC pre-processing pipeline, due to the elimination of a higher number of ICs in the latter.

### 2.7. Statistical Analysis

Differences in the macrostructure metrics between the two nights were calculated by comparing the total time spent in bed, the total sleep time, the wake after sleep onset (i.e. total duration of awake periods from sleep onset), and the percentage of sleep time spent in N2, N3 and REM sleep. Comparisons between the two nights were based on paired t-tests (p<0.05, two-tailed). Statistical differences in the number of rejected ICs and their explained variance between vigilance states were assessed using Friedman test followed by Wilcoxon signed-rank tests (p<0.05) with Bonferroni correction for the six comparisons performed across states.

Non-parametric cluster-based permutation statistics (p<0.05, two-tailed; 5000 permutations) were employed to examine differences in evoked activity obtained from the two pre-processing pipelines. We clustered individual data samples within the −100 ms to 600 ms relative to the event of interest showing significant t-values (p<0.05, two-tailed) based on the temporal and spatial proximity at the sensor level. Each cluster was assigned to cluster-level statistics corresponding to the sum of the t-values of the samples belonging to that cluster. We then evaluated the maximum cluster-level statistics by randomly shuffling condition labels 5000 times to estimate the distribution of maximal cluster-level statistics obtained by chance and applied a two-tailed Monte-Carlo p-value. To assess the magnitude of significant differences, effect sizes were computed using Cohen’s d at the latencies where the largest number of electrodes showed significant differences following permutation testing.

Using the same cluster-based permutation statistical analysis approach, we compared the average ECG signals obtained from the same trials used to compute the HEPs in the two pre-processing pipelines after segmenting the continuous ECG signals from –100 ms to +600 ms relative to the R-peaks during the baseline. This control analysis was carried out to ensure that the observed HEP differences between the standard and PPC pre-processing pipelines did not result from variations in the underlying cardiac signals but rather reflected the impact of the pre-processing strategy.

## 3. Results

### 3.1. Sleep Macrostructure and dataset

Out of 33 volunteers, two were excluded following the first adaptation night, a first withdrew from the study for personal reasons and a second was not asked to return for the experimental nights due to poor sleep quality during adaptation. A third participant was excluded from the analysis due to poor sleep quality upon auditory stimulation on the first experimental night, after which he did not return for the second experimental night. Therefore, the final cohort included in this work was comprised of 30 volunteers with one individual excluded from the REM sleep and another from the N3 sleep analyses due to insufficient valid trials.

Paired t-tests revealed no statistically significant differences (p>0.05) in sleep macrostructure between the two experimental nights (Table S1). Data from the two nights were therefore concatenated for upcoming EEG analyses. Across the two nights, for the HEP analysis, the final sample for each pre-processing pipeline dataset included 30 subjects for wakefulness (800 ± 107 trials, mean ± SD, range = 497-978), 30 subjects for N2 (2221 ± 785 trials, range = 712-3934), 29 subjects for N3 (597 ± 332 trials, range = 123-1232), and 29 subjects for REM (974 ± 325 trials, range = 108-1685). For the AEP analysis, 30 subjects were included for wakefulness (860 ± 28 trials, range = 722-883), 30 for N2 (2745 ± 858 trials, range = 718-4156), 29 for N3 (872 ± 419 trials, range = 175-1609), and 29 for REM (1045 ± 470 trials, range = 117-2362).

### 3.2. PPC Threshold results

The PPC thresholds across vigilance states were 0.07 for wakefulness, 0.10 during NREM sleep and 0.11 during REM sleep. Since PPC values range from 0 (no phase-locking) to 1 (perfect phase-locking), these thresholds suggest that the weakest CFA components used as a reference showed progressively stronger phase-locking from wakefulness to NREM and REM sleep.

### 3.3. Features of ICs

In the standard pre-processing pipeline, a single IC containing the CFA throughout the vigilance state’s duration was observed in 3 participants during wakefulness, 2 during N2 sleep, 1 during N3 sleep and 4 participants during REM sleep.

The visual inspection of ICs exhibiting PPC exceeding the threshold in at least one time-window for each vigilance state revealed that most ICs displayed a broad, slow, and prolonged waveform temporally aligned with the QRS complex of the ECG, resembling characteristics of a PA (Figure 1E; Figure S1). While the morphology of the pulse-related signal showed some variability, this pattern was consistently observed in all included participants across vigilance states (Table 1). Visual inspection of the identified components confirmed artifact-related activity in 6.0 ± 2.2 ICs in wakefulness, 13.0 ± 5.8 ICs in N2, 4.4 ± 2.5 ICs in N3 and 7.6 ± 4.5 in REM sleep per participant. While cardiac related activity in these identified components was mostly related to PAs, in rare cases (less than 1% of the total number of components), CFAs were observed.

**Table 1.**
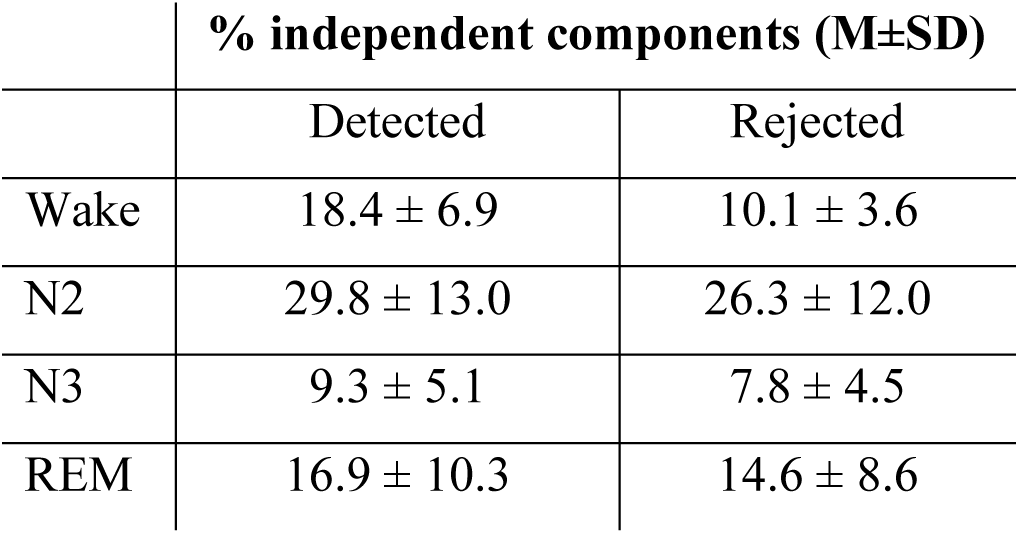
Summary of cardiac-related independent components (ICs), identified in the pairwise phase consistency (PPC) pipeline, detected using the selected PPC threshold and rejected following visual inspection.

The number of rejected cardiac related components, including both CFA and PA, was higher (Bonferroni-corrected Wilcoxon signed rank test, p<0.05) during N2 sleep compared to all other vigilance states, and higher during REM compared to N3 sleep (Figure 2A). In addition, the explained variance of the rejected components was higher in N2 and REM sleep compared to wakefulness and N3 sleep (Figure 2B).

**Figure 2.**
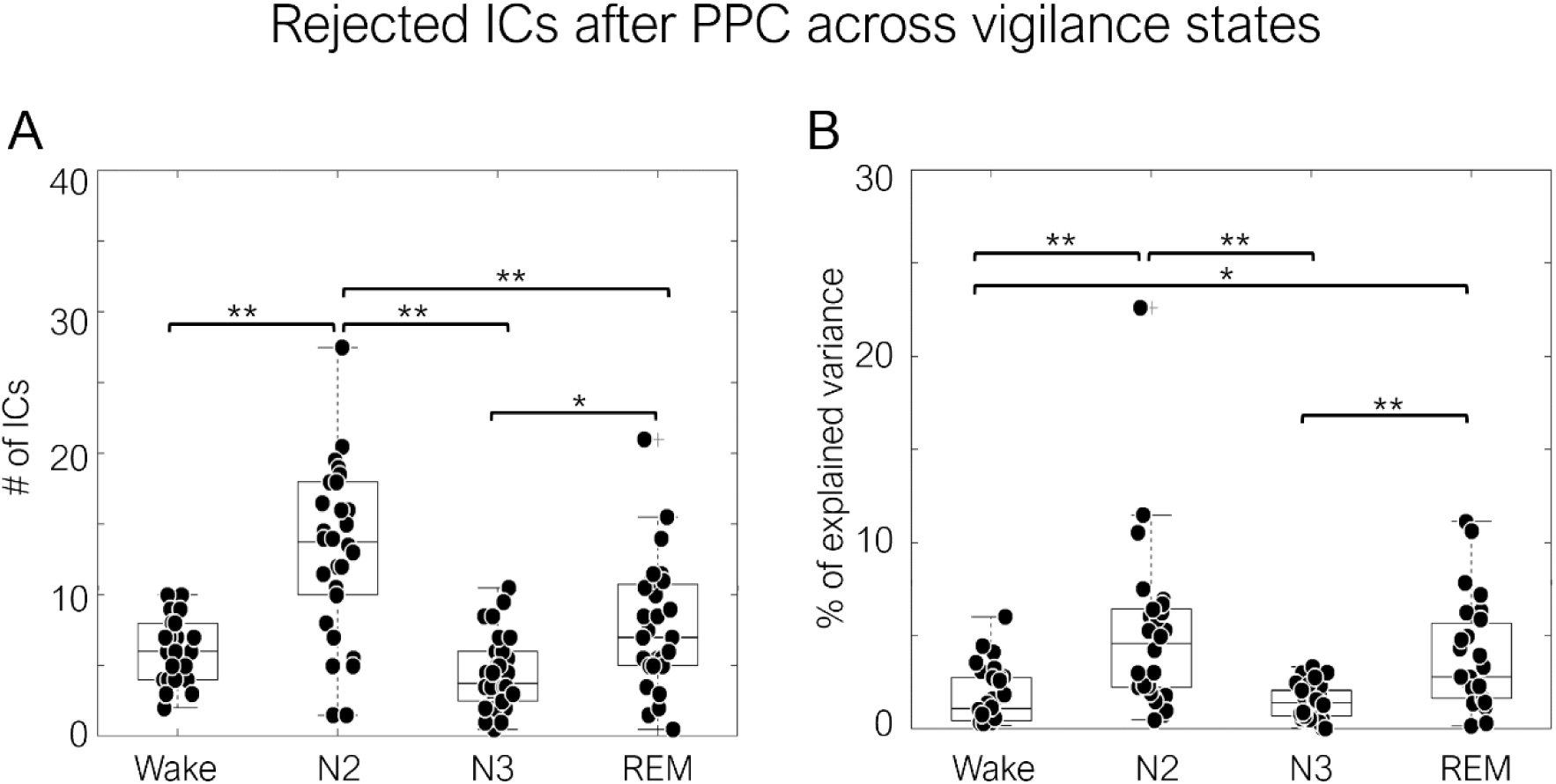
Independent components (ICs) rejected using the PPC pipeline. Number of rejected ICs across participants (A) and their cumulative explained variance as a percentage (B) across vigilance states in wakefulness (N=30), N2 sleep (N=30), N3 sleep (N=29), and REM sleep (N=29). Each black point represents a subject (for sleep stages, data were averaged across the two experimental nights). Boxes display the 25th to 75th percentile, horizontal bars indicate the median, and error bars show the 95% confidence interval. Asterisks indicate statistically significant differences between vigilance states (*p<0.05, **p<0.01 using a Friedman test with post-hoc Wilcoxon signed-rank tests and Bonferroni correction for multiple comparisons).

### 3.4. HEPs

The HEP waveforms showed a prominent early potential time-locked to the R-peak, observed across both pre-processing pipelines and all vigilance states. As expected, given the high-amplitude background activity characteristic of slow-wave sleep, this peak was enhanced in N3 sleep, particularly in its negative deflection [8]. We hypothesized that differences in the HEPs would emerge between the two pipelines, with lower amplitudes for the PPC pre-processing due to the attenuation of cardiac-related artifacts. As a qualitative observation, both the average GFP of the HEP, reflecting overall EEG signal strength, and the absolute HEP magnitude appeared higher in the signal processed with the standard pipeline compared to the PPC pipeline, particularly in wakefulness, N2 and REM sleep. This result was confirmed by statistical analyses comparing HEPs derived from the basic and PPC pipelines within a −100 ms to +600 ms window relative to the R-peak, revealing significant differences across all vigilance states except N3 sleep, where no differences were observed (Figure 3D; p>0.05, two-tailed).

**Figure 3.**
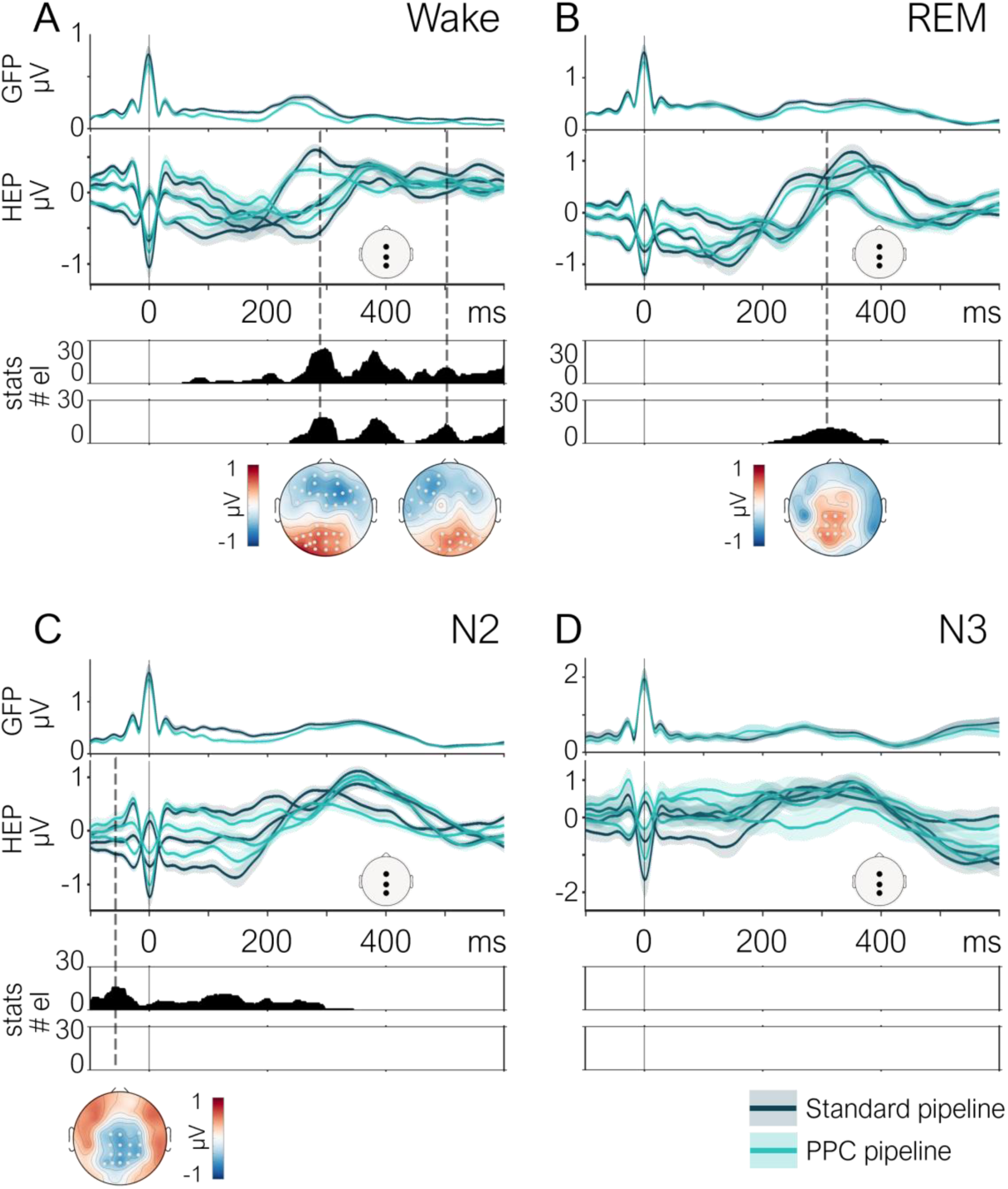
Heartbeat evoked potentials (HEPs) using the standard and PPC pre-processing pipelines in wakefulness and sleep. Grand-average HEP comparison of the standard and PPC pre-processing pipelines across vigilance states in wakefulness (A, N=30), REM sleep (B, N=29), N2 sleep (C, N=30), and N3 sleep (D, N=29). For each vigilance state, the panels show (top to bottom): grand-average GFP; grand-average HEP, displayed for three representative electrodes (Fz, Cz, Pz; locations shown in white topographic schema; dark blue lines for standard pipeline and light blue lines for PPC pipeline); cluster permutation statistical analysis results contrasting HEPs between pipelines shown as the number of significant electrodes (p<0.05, two-tailed), where the upper plot displays negative clusters and the lower plot displays positive clusters of electrodes. The negative/positive clusters include electrodes exhibiting negative/positive and significant differences between the HEPs derived from the two pipelines. The y-axis represents time, with zero corresponding to R-peak latency. Dashed lines indicate the maximum number of electrodes for each significant cluster, with corresponding topographical distributions of HEP differences between pipelines shown at the bottom. White dots on topographic maps indicate significant electrodes.

In the HEP comparison between pre-processing pipelines during wakefulness, the cluster-based permutation test identified four significant clusters (Figure 3A; p<0.05, two-tailed). A first negative cluster spanned approximately 50-435 ms following the R-peak (p < 0.01, Cohen’s d = 1.08), followed by a second negative cluster from 440–600 ms (p = 0.02, Cohen’s d = 0.89). Two positive clusters were also observed, a first between 230–420 ms (p < 0.01, Cohen’s d = 1.92), and a second from 440–600 ms (p = 0.04, Cohen’s d = 0.64). Topographical analysis at the peak of each cluster showed a fronto-central distribution for the negative clusters and a parieto-occipital distribution for the positive clusters. The same analysis conducted for REM sleep revealed a single positive cluster spanning 200–400 ms post-R-peak (p = 0.04, Cohen’s d = 0.95), predominantly localized over parietal scalp electrodes (Figure 3B). In N2 sleep, a single negative cluster was identified between −100–340 ms (p < 0.01, Cohen’s d = 0.88) again, over parietal electrodes (Figure 3C). In addition, we calculated the effect sizes for the observed differences between pipelines. Effect sizes varied by vigilance state, with the strongest effects observed in wakefulness (Cohen’s d up to 1.92), followed by REM (d = 0.95) and N2 (d = 0.88).

Overall, our results indicate that differences between the two pre-processing pipelines were observed irrespective of the degree of vigilance. These differences spanned long intervals, with significant clusters identified as early as ∼50 ms and persisting up to 600 ms after the R-peak, depending on the vigilance state. In wakefulness, four distinct clusters were observed across both early and late latencies, indicating a broad temporal distribution of differences between pre-processing pipelines that extended well beyond the cardiac cycle onset. The absence of significant differences in N3 sleep aligns with our prediction that high amplitude graphoelements would mask cardiac artifacts, making their removal less detectable. Our results show that despite similar threshold values across vigilance states, a higher number of ICs were removed in N2 sleep compared to wakefulness, and the total variance explained by removed components was higher in N2 and REM sleep in comparison to wakefulness. This suggests that the larger differences observed in wakefulness were not simply due to the threshold used for the identification of cardiac-related components.

Control analysis based on the comparison of the ECG between the two pipelines for each vigilance state revealed no significant differences (Figure S2A; p>0.05; two tailed). These results exclude that HEP differences between pre-processing pipelines were trivially explained by differences in the underlying cardiac activity.

### 3.5. AEPs

To evaluate the specificity of the cardiac artifact removal and ensure that neural activity not time-locked to the cardiac cycle was preserved, we compared AEPs across the two pre-processing pipelines. Since AEPs are not time-locked to the cardiac cycle, they serve as a useful control to verify that PPC pre-processing does not distort stimulus-evoked activity. Qualitatively, the characteristic AEP components, including the early N100 observed during wakefulness, as well as the progressive emergence of later components such as the P200 and N550 with increasing sleep depth [12], were identifiable and exhibited comparable amplitudes and latencies across both pre-processing pipelines. Statistical analyses confirmed no significant differences in AEP waveforms between the two pipelines across all vigilance states (Figure 4; p>0.05, two-tailed). These findings suggest that the PPC pre-processing selectively attenuated cardiac artifacts without compromising the integrity of stimulus-related neural activity.

**Figure 4.**
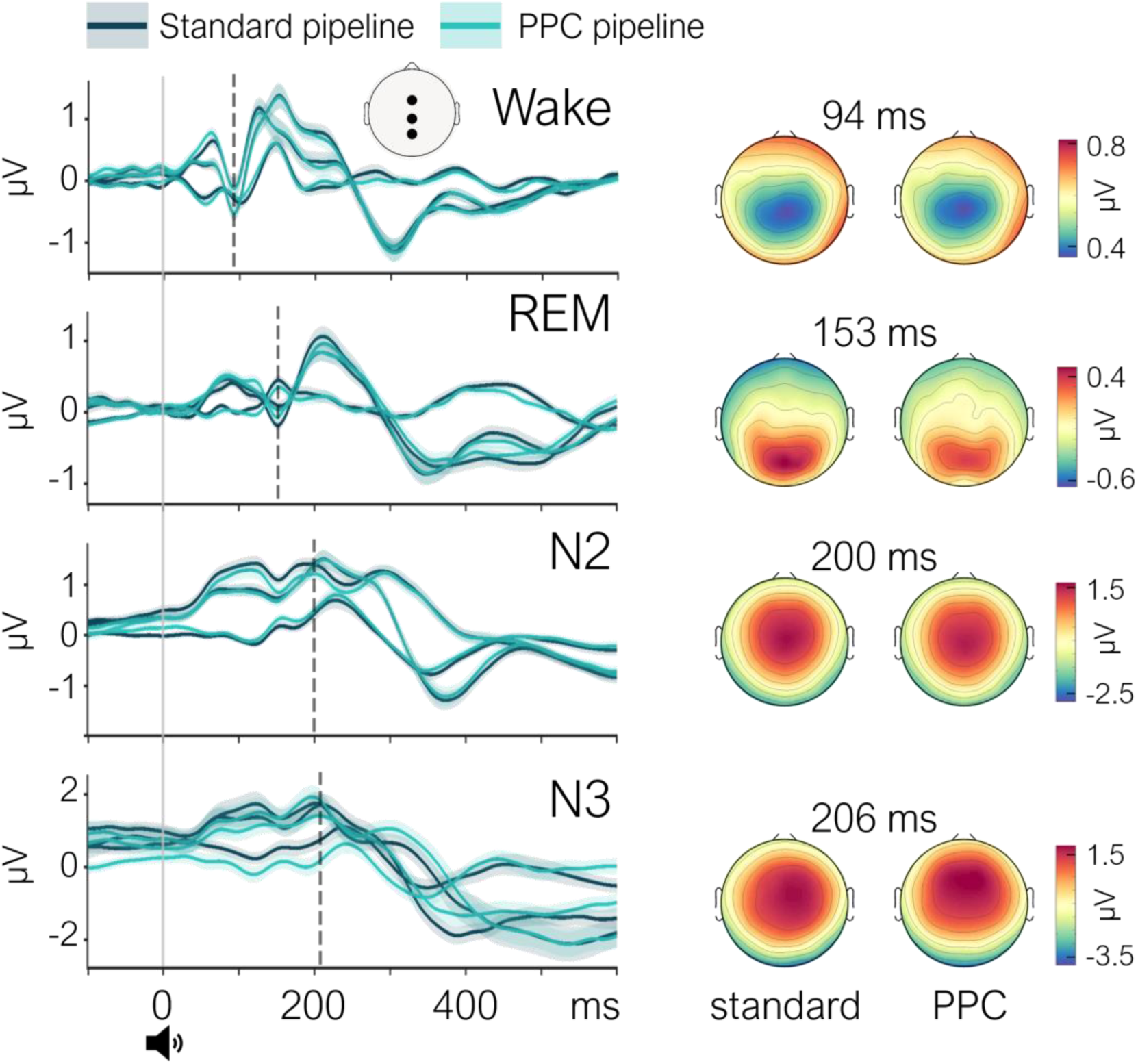
Auditory evoked potentials (AEPs) using the standard and PPC pre-processing pipelines in wakefulness and sleep. Grand-average AEP comparison of standard and PPC pre-processing pipelines across vigilance states. Data are displayed for three representative electrodes (Fz, Cz, Pz; locations shown in white topographic map) across vigilance states (top to bottom): wakefulness (N=30), REM sleep (N=29), N2 sleep (N=30), and N3 sleep (N=29). Dark blue lines for the standard pipeline and light blue lines for the PPC pipeline. Cluster permutation statistical analyses revealed no significant differences between pipelines for any vigilance state (p>0.05, two-tailed). Right panels show topographical distributions of the evoked components for both pipelines at the latencies indicated by dashed lines.

## 4. Discussion

The primary aim of this study was to systematically evaluate the impact of PAs on the HEP, which is inherently time-locked to cardiac activity and therefore particularly susceptible to contamination, and the AEP, a representative example of evoked activity in response to stimuli without temporal alignment to the ongoing heartbeat. We compared the output of two distinct pre-processing approaches: a conventional ICA-based pipeline relying on visual inspection to remove components reflecting CFA (standard pipeline), and a semi-automated method combining ICA with PPC to enhance the detection and removal of cardiac-related components, including those associated with the PA (PPC pipeline). By providing a quantitative, heartbeat-locked synchronization metric, PPC allowed for the systematic and objective identification of candidate pulse-artifact related components, verified through visual inspection. We assessed the impact of these two pipelines on HEPs and AEPs across multiple vigilance states i.e., wakefulness, N2, N3, and REM sleep, to determine whether the impact of PA contamination and the efficacy of artifact correction vary depending on the neural background activity and cardiovascular dynamics (Figure S2B).

### 4.1. Pulse artifacts are consistently observed in wakefulness and sleep

In the standard pre-processing pipeline, ICs reflecting CFA were detected and removed in a small number of participants, ranging from 1 to 4 across vigilance states. As a result, this approach is likely similar to a pre-processing strategy without any cardiac-related artifact correction. In contrast, the PPC pipeline identified and removed additional components in all participants’ datasets, which allowed for a systematic attenuation of PAs. The interpretation of the PPC ICs as pulse-related artifacts is supported by several observations: (i) the ICs removed by the PPC pipeline exhibited smooth, wave-like or saw-tooth morphologies, matching descriptions of PAs [28] (Figure S1); (ii) the differences between pipelines occurred predominantly at later latencies, typically beyond 300 ms post-R-peak, and extended across prolonged intervals (Figure 2), outside the window typically associated with CFA; (iii) these effects were most evident in wakefulness and REM sleep, when the relatively low-amplitude EEG background may allow for subtle cardiac artifacts such as the PA and not the CFA, to be more readily observed. These results demonstrate that cardiac artifact correction, particularly the decision to remove pulse-related contamination, meaningfully shapes the resulting HEP signal.

### 4.2. Heartbeat evoked potential is affected by the pulse artifacts

Our results revealed that the choice of pre-processing pipeline significantly influenced HEP amplitude, particularly in wakefulness, where the differences between the HEPs from the two pre-processing pipelines were sustained over time and extended from 50 ms and up to 600 ms post-R-peak (Figure 3A). Similar but less pronounced effects were observed in REM and N2 sleep: REM exhibited differences between 200 ms to 400 ms after the R-peak (Figure 3B), and N2 showed differences closer to the R-peak between −100 ms to 340 ms relative to the R-peak (Figure 3C). In contrast, HEPs during N3 were not affected by the type of pre-processing, likely due to the high-amplitude EEG background that may mask the expression of PAs (Figure 3D). This state-dependent pattern supports the notion that the prominence of PAs, and thus the impact of their correction, may depend on the underlying EEG dynamics.

Of note, while evoked activity time-locked to the ongoing heartbeat was affected by the removal of cardiac artifacts, evident in the HEP differences between pipelines, evoked activity unrelated to the cardiac rhythm, specifically AEPs, did not show statistically significant differences between pre-processing procedures (Figure 4). This highlights the specificity of the PPC pre-processing approach in isolating and removing cardiac artifact-related activity.

The impact of the PPC pipeline on the HEPs during wakefulness and its gradual decrease with increasing sleep depth was also confirmed by the effect size of the HEP differences ranging from 1.92 in wakefulness, followed by 0.95 in REM sleep and 0.88 in N2 sleep. These findings support our hypothesis that PAs can be better isolated and subsequently excluded during states with lower EEG background amplitude and high variability in cardiac activity and blood pressure (wakefulness and REM), moderately isolated during intermediate states (N2), and least isolated during high-amplitude slow-wave activity and low cardiac variability (Figure S2B) and blood pressure (N3).

When examining the distribution of rejected cardiac-related components across vigilance states, we observed that the number of removed components was significantly higher in N2 compared to all other states, and higher in REM compared to N3 sleep (Figure 2). Consistently, the explained variance of the removed components was also higher in N2 and REM sleep compared to both wakefulness and N3. While this pattern is in line with the larger number of rejected components in these stages, neither the number of rejected ICs nor their explained variance directly accounts for the HEP differences observed across pipelines. In accordance, the most pronounced HEP differences were identified in wakefulness despite fewer cardiac-related components being removed, while N2, which showed the highest number of rejected components, revealed only modest HEP differences in terms of effect size and length of significant intervals.

The impact of PA removal in HEP analyses has important methodological implications. In practice, many studies may not adequately address the PA, which can be present even when no CFA components are identified. Some studies attempted to control for cardiac contamination by comparing ECG signals across conditions or populations, reasoning that in the absence of ECG differences, the effect of CFAs is counterbalanced across conditions [13, 50–53]. While this approach may help exclude CFA-related differences, it does not address the PA, which is not an immediate consequence of the cardiac electromagnetic activity reflected in the ECG signal. Other studies addressed this issue by restricting analyses to time windows that are presumed to be free of cardiac contamination [2, 9, 11, 16, 20, 29, 54–61], typically avoiding periods immediately surrounding the R-peak or T-wave. However, our findings challenge this assumption by demonstrating that later time windows, especially those beyond 300 ms post-R-peak, can also be affected by pulse-related activity, especially during wakefulness. This suggests that simply avoiding early latencies is not sufficient to mitigate the impact of cardiac artifacts on EEG signals, and that pulse contamination can affect HEP estimation along the whole time-period after the R-peak up to 600 ms post R-peak onset.

### 4.3. Limitations

We acknowledge that the efficacy of the ICA decomposition during each sleep stage may be limited by its application over long recordings resulting from the concatenation data acquired over time periods spanning several hours. This is especially the case for N2 sleep, which makes up ∼50% of a healthy night of sleep, and even for sleep stages of shorter duration (REM or N3), in all of which discontinuous recording periods from evening to morning are merged. Concatenating these periods of homogenous sleep stages has the advantage of maximizing the information from all the available data and capturing the consistent features across several hours of a recording. However, it also has the disadvantage of posing a limit on the assumption of the semi-stationarity of the EEG data. Another possible factor potentially affecting PA estimation was the pressure on specific electrodes upon EEG acquisition, especially during sleep recordings, depending on the participant’s position in bed. As this position would likely change during an overnight sleep recording, the assumption of the semi-stationarity of the EEG signals prior to the ICA decomposition may be of limited applicability in this context.

### 4.4. Conclusion

Cardiac-related artifacts, including cardiac electromagnetic activity and pulse activity (mainly originating from blood vessel pulsations), are detectable in a variety of vigilance states during wakefulness and sleep. The impact of PA contamination on evoked activity is evident in HEPs from all vigilance states (wakefulness, N2 and REM sleep) except N3, while it remains negligible in AEPs. These findings highlight that PAs should be considered as a significant confounding factor, especially when comparing HEPs between groups that may vary in terms of blood pressure or vascular dynamics.

## Data and code availability

Data and analysis code can be made available by the corresponding author upon reasonable request. The code for the experimental paradigm administration (https://github.com/fcbg-platforms/eeg-cardio-audio-sleep/tree/maint/0.3) is publicly available.

## Author contributions

**Jacinthe Cataldi:** Conceptualization, Data curation, Methodology, Formal analysis, Writing - original draft; **Andria Pelentritou:** Conceptualization, Data acquisition, Data curation, Methodology, Formal analysis, Writing - original draft; **Sophie Schwartz:** Conceptualization, Resources, Funding acquisition, Project administration, Writing - review and editing; **Marzia De Lucia:** Conceptualization, Methodology, Funding acquisition, Project administration, Supervision, Writing - original draft.

## Acknowledgements

We are grateful to Matthieu Scheltienne from the Campus Biotech Foundation in Geneva for designing the code for the experimental paradigm administration. Further, we thank Gwenael Birot and Virginie Sterpenich from the Campus Biotech Foundation in Geneva for useful feedback and guidance with regards to experimental procedures. Finally, we extend our gratitude to Alice Clerget, Erin Mahan and Marie Zocca for assistance with participant recruitment and data acquisition.

