## Supplementary Materials for "Impact of the pulse artifact on evoked activity in human wakefulness and sleep"

**Table S1. Sleep macrostructure across healthy volunteers and experimental nights.** Time in bed calculated based on lights off and lights on times. Total sleep time computed as the period between the sleep onset and final awakening. Percentage of occurrence of each sleep stage (N2, N3, REM) during the session were calculated relative to the total sleep time. Percentage sleep Efficiency was computed as the total sleep time over the time in bed. SD = Standard Deviation.

| | **Mean ± SD** |
|---|---|
| Time in bed (mins) | 499.09 ± 40.83 |
| Total sleep time (mins) | 441.20 ± 44.02 |
| Wake after sleep onset (mins) | 42.85 ± 28.70 |
| Sleep latency (mins) | 4.55 ± 10.20 |
| Sleep efficiency (%) | 88.51 ± 6.90 |
| N2 sleep (%) | 53.55 ± 5.95 |
| N3 sleep (%) | 14.97 ± 7.27 |
| REM sleep (%) | 23.53 ± 4.10 |

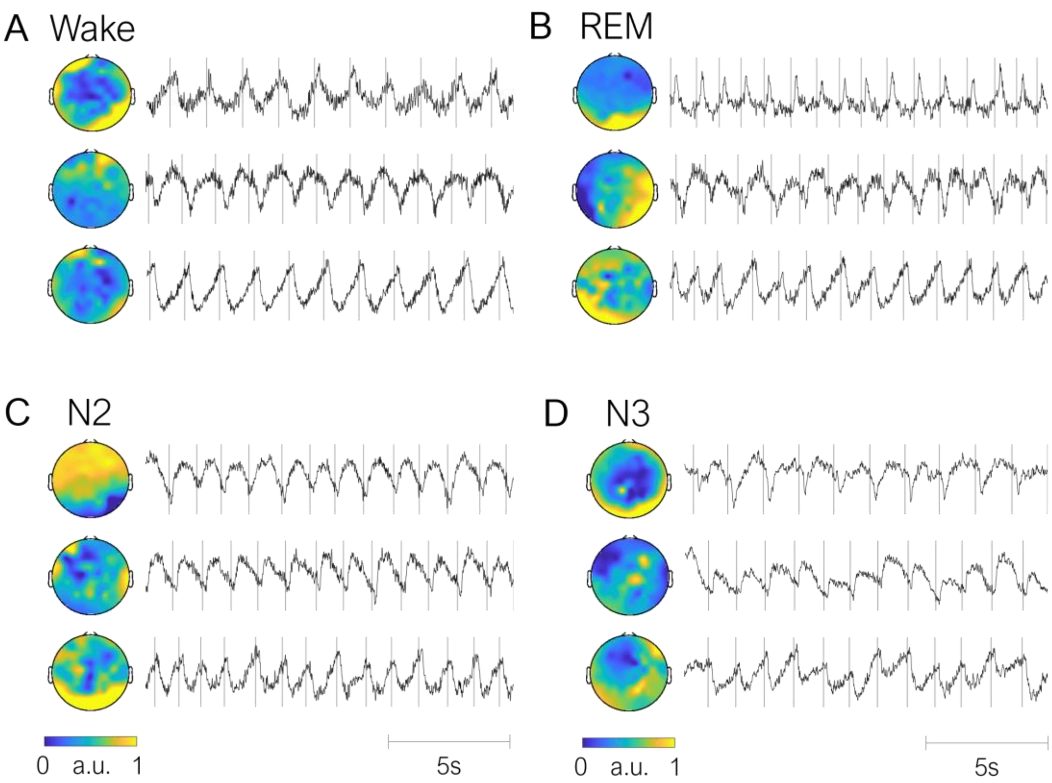


**Figure S1. Pulse artifacts across vigilance states.** Examples of independent components (ICs) displaying pulse artifact (PA) features across vigilance states: wakefulness (A), REM sleep (B), N2 sleep (C), and N3 sleep (D). Each panel shows three example ICs with pairwise phase consistency exceeding the PPC threshold and confirmed as PAs upon visual inspection. For each IC, the topographic distribution averaged over the entire component (left) and 20-second time course (right) are displayed. Vertical gray lines indicate R-peak timing. The slow-wave activity consistently follows the R-peaks in a systematic, time-locked manner, which is the primary feature characteristic PAs. This temporal relationship is particularly evident when there are variations in heart rate with corresponding changes in the timing of PA waves (e.g., the third IC in panel B).

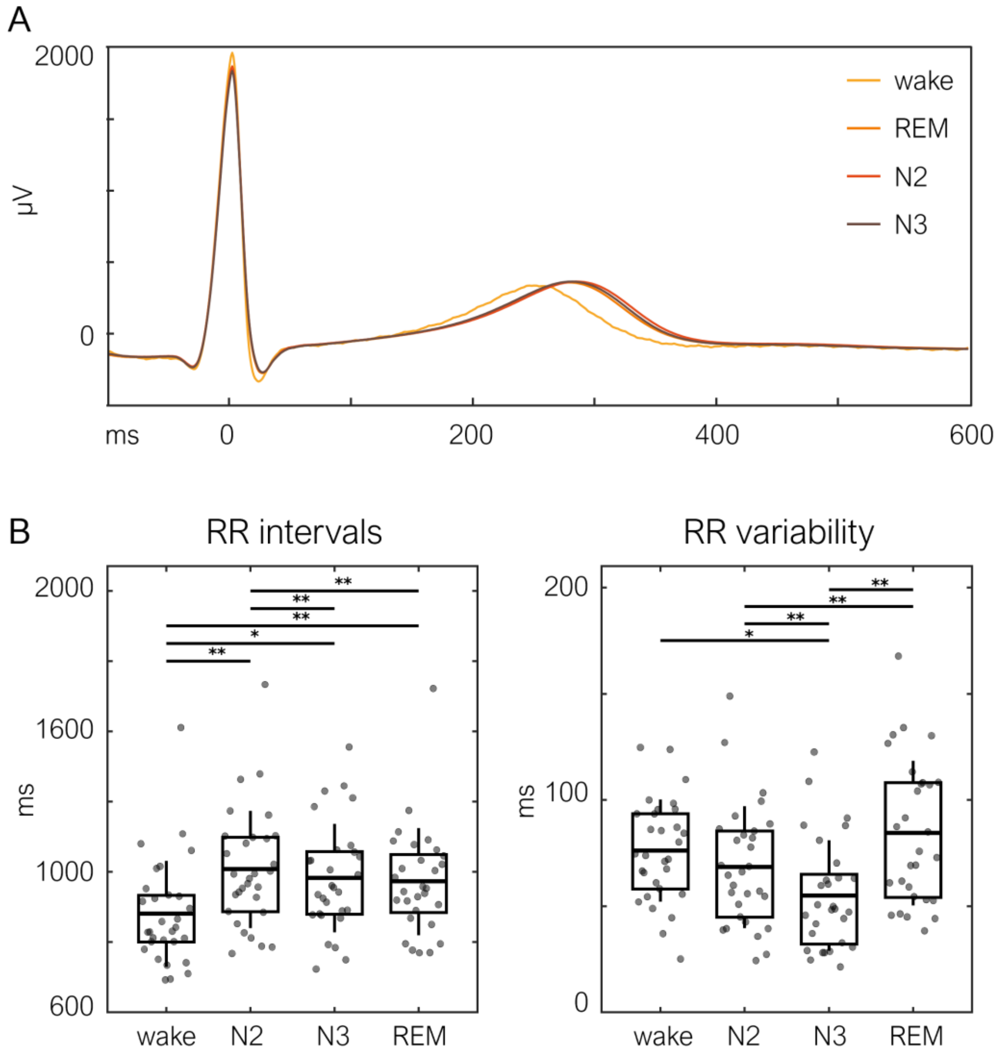


**Figure S2. ECG characteristics across vigilance states.** A. Grand average ECG waveforms across trials selected for the PPC pre-processing pipeline, shown for each vigilance state: wakefulness (N=30), REM sleep (N=29), N2 sleep (N=30), and N3 sleep (N=29). Matched waveforms were obtained for the standard pre-processing pipeline. Zero corresponds to the R-peak latency. **B.** Boxplots of R-peak to R-peak (RR) intervals and RR interval variability (standard deviation of RR intervals). Each black point represents a subject (for sleep stages, data were averaged across the two experimental nights). Boxes display the 25th to 75th percentile, horizontal bars indicate the median, and error bars show the 95% confidence interval. Asterisks indicate statistically significant differences between vigilance states (*$p<0.05$, **$p<0.01$ using a Friedman test with post-hoc Wilcoxon signed-rank tests and Bonferroni correction for multiple comparisons).
